# Rapid detection of cytochrome *cd1*-containing nitrite reductase encoding gene *nirS* with loop-mediated isothermal amplification assay

**DOI:** 10.1101/404392

**Authors:** Qianqian Yang, Xuzhi Zhang, Xiaoyu Jiang, Xiaochun Wang, Yang Li, Jun Zhao, Zhihui Hao, Pingping Wang, Keming Qu

## Abstract

The cytochrome *cd1*-containing nitrite reductase, *nirS*, plays an important role in biological denitrification. Consequently, investigating the presence and abundance of *nirS* is a commonly used approach to understand the distribution and potential activity of denitrifying bacteria, in addition to denitrifier communities. Herein, a new molecular biology technique termed loop-mediated isothermal amplification (LAMP) was developed to rapidly detect *nirS* gene using those of *Pseudomonas aeruginosa* to optimize the assay. The LAMP assay relied on a set of four primers that were designed to recognize six target sequence sites, resulting in high target specificity. The specificity of the assay was confirmed by the lack of amplification when using DNA from 15 other bacterial species lacking *nirS* gene. The limit of detection for the LAMP assay under optimized conditions was 1.87 pg/reaction of genomic DNA, which was an order of magnitude lower than that required by conventional PCR assays. Moreover, a cell-template based LAMP assay was also developed for detecting *nirS* gene that directly used bacterial cells as template rather than genomic DNA. Only 1 h was needed from the addition of bacterial cells to the reaction to the verification of amplification success, and bulky and sophisticated equipment were not needed. Further, the *nirS* gene of *P. aeruginosa* in spiked seawater samples could be detected with both the DNA-template based LAMP assay and the cell-template based LAMP assay, thereby demonstrating the practicality of in-field use of them. In summary, the LAMP assays described here represent a rapid, user-friendly, and cost-effective alternative to conventional PCR.

## 1. Introduction

Denitrification that involves the reduction of nitrate to gaseous forms is a globally important process with relevance to many environments (1-3). For example, denitrification can lead to the loss of nitrogen content in agricultural soils, but is also employed to remove excess nitrogen in environments like wastewaters and sludges (2). Microorganism-mediated activities play an important role in denitrification and have even been applied to alleviate eutrophication (1,4,5). Thus, a more detailed understanding of denitrifying organisms will aid in the application of numerous denitrification-related processes. Denitrifying bacteria comprise a wide diversity of microbial species. Cultivation-independent investigation of denitrifiers has been commonly used and has focused on analyzing key reductase functional genes (2-6). In particular, the key step in denitrification is the reduction of nitrite to nitric oxide that is catalyzed by two structurally different, but functionally equivalent, forms of nitrite reductase encoded by the *nirK* and *nirS* genes (2,3,7). Thus, *nir* genes are commonly used molecular markers for characterizing the diversity and abundance of denitrifying bacteria in environmental communities(3,7-9). Of these, *nirS* is frequently used because its phylogenetic signal is largely congruent with that of 16S rRNA genes at the family or genus levels (10,11).

The application of modern molecular biological techniques has greatly facilitated the detection of specific genes. In the last few decades, numerous methods including polymerase chain reaction (PCR) (2,3,11-15), denaturing gradient gel electrophoresis(2,16) and gene chips (17) have been used to detect and analyse *nirS* gene prevalence and diversity. Among these, PCR-based methods have been prominently employed due to their high degree of accuracy and reliability. In particular, quantitative real-time PCR (qPCR) is a highly sensitive and popular tool for *nirS* detection that allows simultaneous quantification (11). However, qPCR suffers from several drawbacks including the requirement of specialized equipment, trained operators, and high costs that largely limit its application in resource-limited settings and, especially, in-filed applications (18,19).

Loop-mediated isothermal amplification (LAMP) that was established by Notomi *et al.* in 2000 (20) has the potential to overcome drawbacks associated with conventional PCR and revolutionize molecular biology. Compared to conventional PCR methods, it exhibits several significant advantages (18) including: 1) Specialized equipment is not necessary due to the avoidance of thermal cycling, resulting in advantages including ease of miniaturization, low energy consumption, and high efficiency (20,21). 2) Higher specificity by LAMP is achieved due to the use of four to six different primers that bind specific sites on the template strand. 3) Sensitivity is less affected by substances that usually inhibit PCR reactions (21,22). These advantages suggest that simple assays could be developed using LAMP with elimination of the most cumbersome steps of sample pretreatment including DNA extraction and purification (23-25). Several studies have demonstrated the capacity of LAMP to directly amplify target genes from rapidly processed, crude sample matrices (26-29) including unprocessed samples with or without simple mechanical-based pretreatments (25,30,31). Consequently, the employment of LAMP considerably reduces the cost and turnaround time associated with gene detection. However, there have been no reports of *nirS* gene detection via LAMP.

We have successfully used LAMP assays previously to detect *malB* genes of *Escherichia coli* (32,33). Herein, we developed a rapid, easy-to-use, and cost-effective approach for realizing in-field detection of *nirS* gene of *Pseudmonas aeruginosa* (34), by constructing a DNA-template based LAMP assay and a cell-template based LAMP assay. The sensitivity and specificity of the new approach were characterized and compared to conventional PCR methods via a sensitivity analysis with extracted genomic DNA as template. Moreover, the LAMP assays were also used to detect *nirS* gene in seawater samples spiked with genomic DNA or *P. aeruginosa* cells.

## 2. Materials and Methods

### 2.1 Bacterial strains

Standard bacterial strains of *P. aeruginosa* (PAO1, ATCC 15692), *E. coli* (ATCC 35150), *Staphylococcus aureus* (ATCC25923), *Listeria monocytogenes* (ATCC 19116), *Salmonella typhimurium* (ATCC 14028), *Vibrio parahaemolyticus* (ATCC 17802), and *Shigella flexneri* (CGMCC11868) were all purchased from BIOBW Biotechnology Co., Ltd (Beijing, China). Additional strains including *E. coli* (120303502, 120303510, 120303512) and *Streptomyces* (1203EC1070400021, 1203SPL070400003, SAHL070400003) were isolated and identified by Zhihui Hao and Pingping Wang from environmental samples taken from farms. *Halomonas alkaliphila* strains (strains X1, X2, X3) were also isolated and identified by Yan Zhang from seawater samples.

### 2.2 Cultivation and cell quantification

Luria-Bertani (LB) medium was used to culture *P. aeruginosa*, *E. coli*, *S. aureus*, *L. monocytogenes*, *S. typhimurium*, *Streptomyces* spp. and *S. flexneri*, while 2216E medium was used to culture *V. parahaemolyticus* and *H. alkaliphila*. Culture media was purchased from the Hope Bio-Technology Co., Ltd (Qingdao, China). Bacterial cultivation was conducted in accordance with previously described methods (35,36) with minor modifications. Briefly, strains were stored at −80°C and then pre-grown overnight in the appropriate medium with constant shaking. The incubation temperature was 37°C unless otherwise indicated. Active strains were then further transferred to new culture medium. After a second incubation for ∼10 h, cell numbers were measured using a plate counting method that we have used previously (36). The cultures were then immediately diluted to achieve the desired cell concentrations for further use, or otherwise centrifuged to collect cells for DNA extraction.

### 2.3 Genomic DNA extraction and purification

DNA was extracted from cells collected from 50 mL of sub-cultured medium, followed by DNA purification using previously described methods (33). Briefly, cells were pre-separated by centrifugation and genomic DNA was extracted and purified from the collected cells using a rapid commercial genomic DNA extraction kit (Biomed Co., Beijing, China) according to the manufacturer’s instructions. Purified DNA was then quantified using a Biodropsis BD-2000 spectrophotometer (Biodropsis Technologies Co., Ltd, Beijing, China). Template genomic DNAs were then stored in Tris– EDTA buffer (pH 7.0) at −20°C until further use no later than four weeks after extraction.

### 2.4 LAMP assays

#### 2.4.1 Primer design and synthesis

The *nirS* gene sequence of *P. aeruginosa* was obtained from the NCBI database (http://www.ncbi.nlm.nih.gov/gene/882217). LAMP primer sets to amplify *nirS* were designed based on the published DNA sequence using the LAMP primer designing software package (v.4.0, https://primerexplorer.jp/e/). Using previously published guidelines (37), the specificity of the primers was determined using the NCBI BLAST (Basic Local Alignment Search Tool), and then screened using Primer Premier v.5.0 (PREMIER Biosoft International, Palo Alto, CA) based on the likelihood of primer dimer formation and non-specific priming. From these analyses, a single primer set was selected for LAMP assays (Figure 1). From the first bate of F3 to the last bate of B3, there were 207 bp. The sequence of the 207 bases was checked using the NCBI BLAST. Only *nirS* gene of a few *P. aeruginosa* strains matched at 100%. The primers were then synthesized by Sangon Biotech Co., Ltd, (Shanghai, China). The priming locations on the target DNA sequence are shown in Figure 1, and the primer nucleotide sequences are provided in Table 1.

**Table 1.**
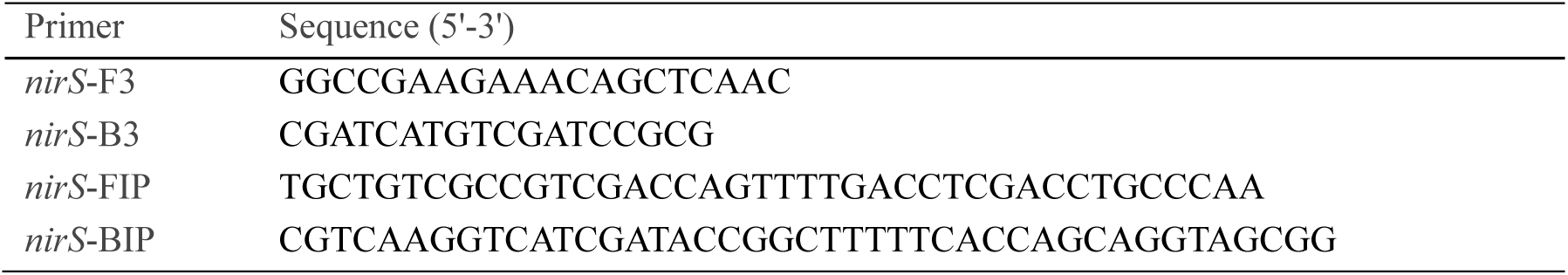
LAMP primer sequences.

**Figure 1.**
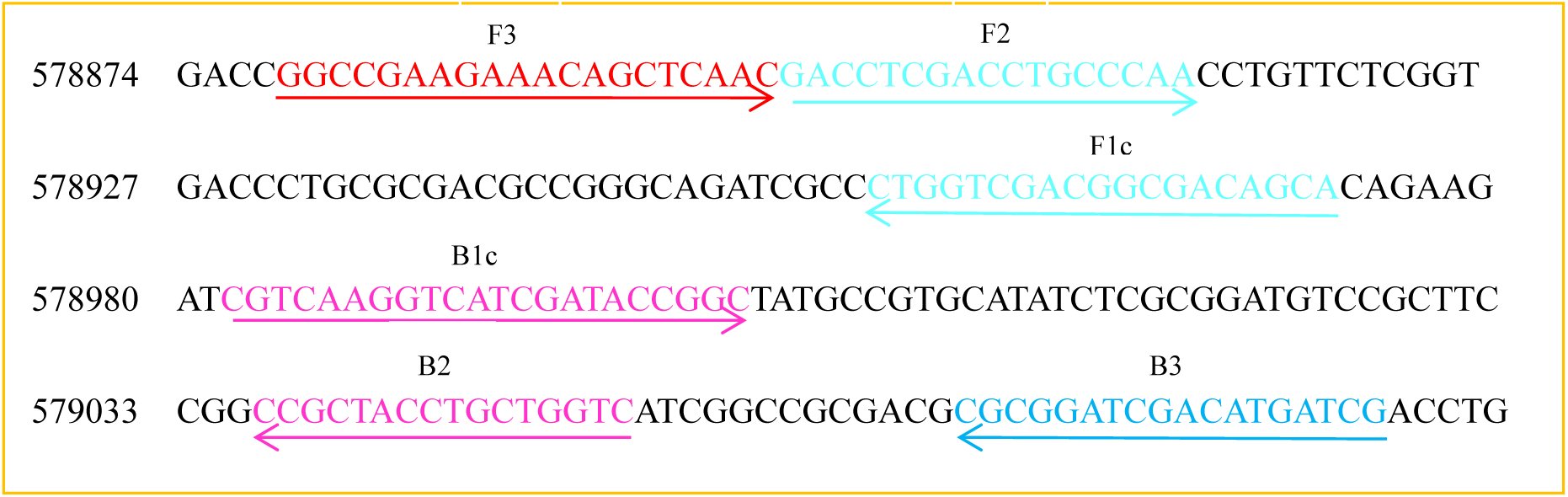
Priming locations and orientation of the LAMP primers developed to amplify *P. aeruginosa nirS*. The arrows show sequence directions from 5’ to 3’. The asterisks denote consistent nucleotides sequence not shown.

#### 2.4.2 LAMP reaction systems and amplification product characterization

As shown in Scheme 1a, LAMP assays using DNA as template, termed DNA-template based LAMP assays, were conducted using previously described methods (20,33) with minor modifications. Unless otherwise indicated, 25 μL LAMP reaction volumes were used comprising 0.2 μM of each outer primer (B3 and F3), 1.6 μM of each inner primer (FIP and BIP), 1.2 mM of each dNTP, 2.5 μL 10× ThermoPol^®^ reaction buffer, 1 μL *Bst* 2.0 DNA polymerase, 6 mM MgSO_4_, and 1 μL genomic DNA template. dNTPs were purchased from MBI Fermentas (Waltham, USA) and *Bst* 2.0 DNA polymerase was purchased from New England Biolabs (Ipswich, Massachusetts, USA). Reactions were incubated at 63°C for 60 min in a block heater, unless otherwise indicated. Based on the methods described in Tomita *et al.* (38), LAMP reaction products were characterized by gel electrophoresis on a 2% agarose gel using a DY-6 electrophoresis apparatus (Xinghua Assay Apparatus Factory, Beijing, China) and a DNR Bio-Imaging System (MF-ChemiBis 3.2, Israel). Electrophoresis was conducted using 50x diluted LAMP reaction products. Additional visual characterization using the fluorescent dye GeneFinder (Biov Co., Ltd., Xiamen, China) was also performed. Triplicate determinations were performed for every set of analyses.

As shown in Scheme 1b, cell-template based LAMP assays were carried out using the same method described for the *nirS* gene, but with *P. aeruginosa* cells as template rather than extracted genomic DNA. *P. aeruginosa* cells were obtained using the methods described by Kanitkar *et al.* (30). Briefly, after the concentration of bacterial cells was quantified using the plate counting method described above, 10 mL of culture medium was centrifuged at 13,000 g for 15 min to obtain a biomass pellet. The biomass pellet was then suspended in an appropriate volume of water and 2 μL of the bacterial suspension was immediately used as amplification template. LAMP products were again characterized by gel electrophoresis and fluorescent dye visualization as described above.

#### 2.4.3 Optimization

The temperatures and incubation times for the LAMP assay were optimized based on the approach of Balbin *et al.* (39). Briefly, amounts of *P. aeruginosa* genomic DNA varying from 18.70 fg–187.00 ng were used as amplification template. LAMP was then carried out at 61°C, 62°C, 63°C, 64°C, and 65°C. After determining the optimal temperature for the assays, LAMP was then conducted with varying incubation times including 10, 20, 30, 40, 50, 60, 70, and 80 min.

### 2.5 Specificity

The specificity of the designed *nirS* primer set was determined using either genomic DNA or bacterial cells as amplification templates. For the former, ∼0.1 ng genomic DNA from *P. aeruginosa*, *E. coli*, *S. aureus*, *L. monocytogenes*, *S. typhimurium*, *Streptomyces* spp., *S. flexneri*, *V. parahaemolyticus*, *H. alkaliphila X1*, *H. alkaliphila X2* or *H. alkaliphila X3* was used as template for the LAMP assay. Gel electrophoresis and visual detection were used to characterize the amplification products. For assays with cells, ∼10^5^ CFU of cells were used as amplification template. For both sets of assays, 0.19 ng of *P. aeruginosa* genomic DNA and pure water were used as the positive and negative controls, respectively.

### 2.6 Sensitivity

#### 2.6.1 Sensitivity of DNA-template based LAMP assay

The sensitivity of the DNA-template based LAMP assay for *nirS* was determined using a 10-fold serial dilution of the template DNA. Results were again characterized using both gel electrophoresis and visual detection. The limits of detection (LOD) were obtained from these analyses using the same reaction parameters discussed above. Unless otherwise indicated, each assay was conducted in triplicate.

#### 2.6.2 Sensitivity of cell-template based LAMP assay

The sensitivity of the cell-template based LAMP assay for *nirS* was determined using methods described by Lee *et al.* (26) with minor modifications. Briefly, a biomass pellet of bacterial cells was obtained from centrifugation of the cell cultures. The pellet was then suspended in 5 mL water. An aliquot (500 μL) of the bacterial suspension was used to measure cellular concentrations. The remainder of the suspension was used as template for direct amplification using the LAMP assay with 10-fold serial dilutions to identify the LOD (CFU/reaction). The results were characterized with both gel electrophoresis and visual detection. Unless otherwise indicated, each assay was conducted in triplicate.

### 2.7 Conventional PCR assays

The F3 and B3 primers were used for conventional PCR assays following the methods of Verma *et al.* (19) and Stedtfeld *et al.* (31). PCR reactions comprised 25 μL volumes consisting of 1 μL genomic DNA template, 12.5 μL Version 2.0 Taq polymerase plus dye (TaKaRa Biotechnology Co., Ltd., Dalian, China), and 1 μL of each primer (0.2 μM in reaction mix). PCR reaction conditions consisted of 94°C for 3 min, followed by 30 cycles of 94°C for 45 s, 54°C for 55 s, 72°C for 90 s and a final extension at 72°C for 10 min. A 5 μL aliquot of each PCR product was subjected to 2% agarose gel electrophoresis for characterization.

### 2.8 Detection of nirS gene in spiked seaweater samples

To investigate the ability of the LAMP assay to detect *nirS* in complex natural matrices, a spiked LAMP assay was conducted with seaweater samples. The seawater sample was collected from the Yellow Sea (36°06.54′;N; 120°39.28′E). Water salinity (31.01‰) and pH (8.062) were measured using a YSI 556 Multi Probe System (Envisupply Co., USA). Bacterial biomass and extracellular DNA were removed from the water using filtration with 0.22 μm Sterivex filters followed by filtration with Silicone membranes (EMD Millipore Corp., Billerica, MA), respectively (31). The capacity of the LAMP assays to detect *nirS* gene was then challenged using seawater samples spiked with genomic DNA and *P. aeruginosa* cells, respectively. All seawater samples were used for the next step within 20 min after the spiked performance, unless otherwise indicated.

#### 2.8.1 DNA-template based LAMP assay

Extracted *P. aeruginosa* genomic DNA was added to the filtered seawater over a concentration range of 1.27 × 10^2^–1.27 × 10^8^ fg/μL. Then, 1 μL of seawater samples with varying spiked DNA concentrations were directly used as templates for LAMP assays. The LOD were then determined based on visual detection of the amplification products.

#### 2.8.2 Cell-template based LAMP assay

A ∼10^13^ CFU/mL bacterial suspension was prepared in water, as described above. Bacterial suspensions were added to the filtered seawater at various volumes to generate spiked samples over a cell concentration range of 1.68 × 10^1^–1.68 × 10^7^ CFU/mL. For each spiked sample, a 50 mL cell suspension was subjected to centrifugation to pellet cells. The obtained biomass pellet was then directly used as template for LAMP assays. In addition, 2 μL of spiked seawater samples were directly used as templates for LAMP assays. The LOD were determined based on visual detection of amplification products.

## 3. Results

### 3.1 LAMP amplification of *nirS*

Using 0.19 ng genomic DNA of *P. aeruginosa* as template, LAMP amplification of *nirS* at 63°C for 60 min resulted in amplification products of various size, as indicated by gel electrophoresis and the presence of many sized bands in a reproducible ladder-like pattern(Figure 2a), which was the same phenomena obtained somewhere (20-23,32). The absence of these ladder-like patterns in the negative control indicated that the pattern was due to *nirS* amplification. Light green fluorescence of positive amplification products when using the GeneFinder dye was evident (Figure 2b) as previously observed (32), while negative controls remained orange. To avoid inhibition of the dye fluorescence, 1 μL of GeneFinder solution was coated inside of the Eppendorf tube cover, rather than premixing it into the LAMP reaction mixture.

**Figure 2.**
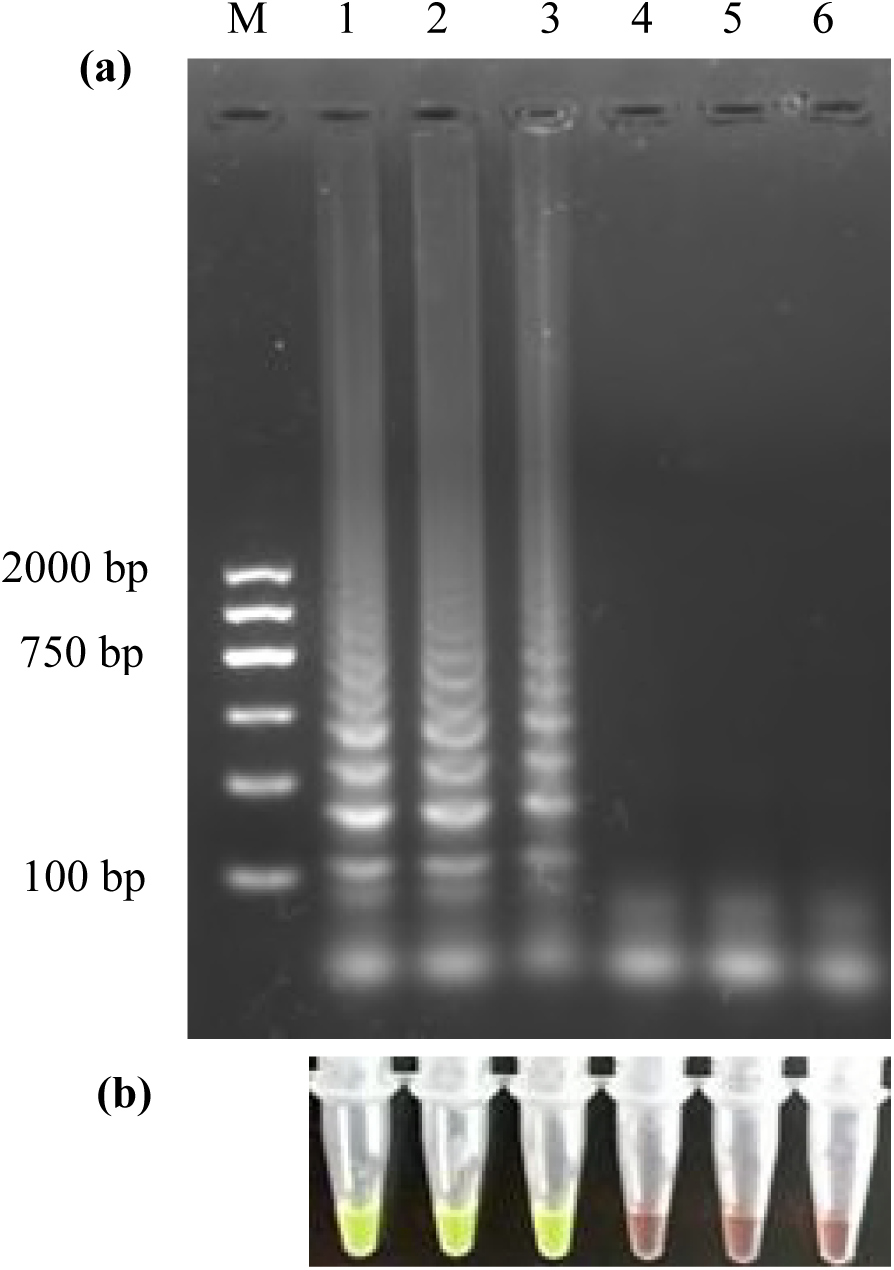
Detection of the *P. aeruginosa nirS* gene with triplicate LAMP assays using genomic DNA as templates. Amplification success was verified by gel electrophoresis (a) and GeneFinder (b). Lanes 1–3 are amplification reactions using 0.19 ng of extracted genomic DNA as template; Lanes 4–6 are the negative control (using sterile water as template). The LAMP reactions were incubated at 63°C for 60 min.

To optimize the assay, LAMP reactions were conducted at various temperatures and incubation times. The ladder-like electrophoresis patterns observed in the initial amplifications were reproduced when using 0.19 pg of genomic DNA as template and incubating reactions at 63°C for 60 min. Modifying the incubation temperatures or using incubation times < 60 min resulted in the absence of ladder-like electrophoresis band patterns. Consequently, an incubation temperature of 63°C and time of 60 min were selected for further analyses.

Using 3.36 × 10^2^ CFU of *P. aeruginosa* cells as template, cell-template based LAMP assays were also incubated at 63°C for 60 min and yielded similar successful amplification results as with amplification using genomic DNA (Figure 3a and 3b), without negative control amplification. These results indicated positive LAMP amplification from *P. aeruginosa* cells under the specified conditions.

**Figure 3.**
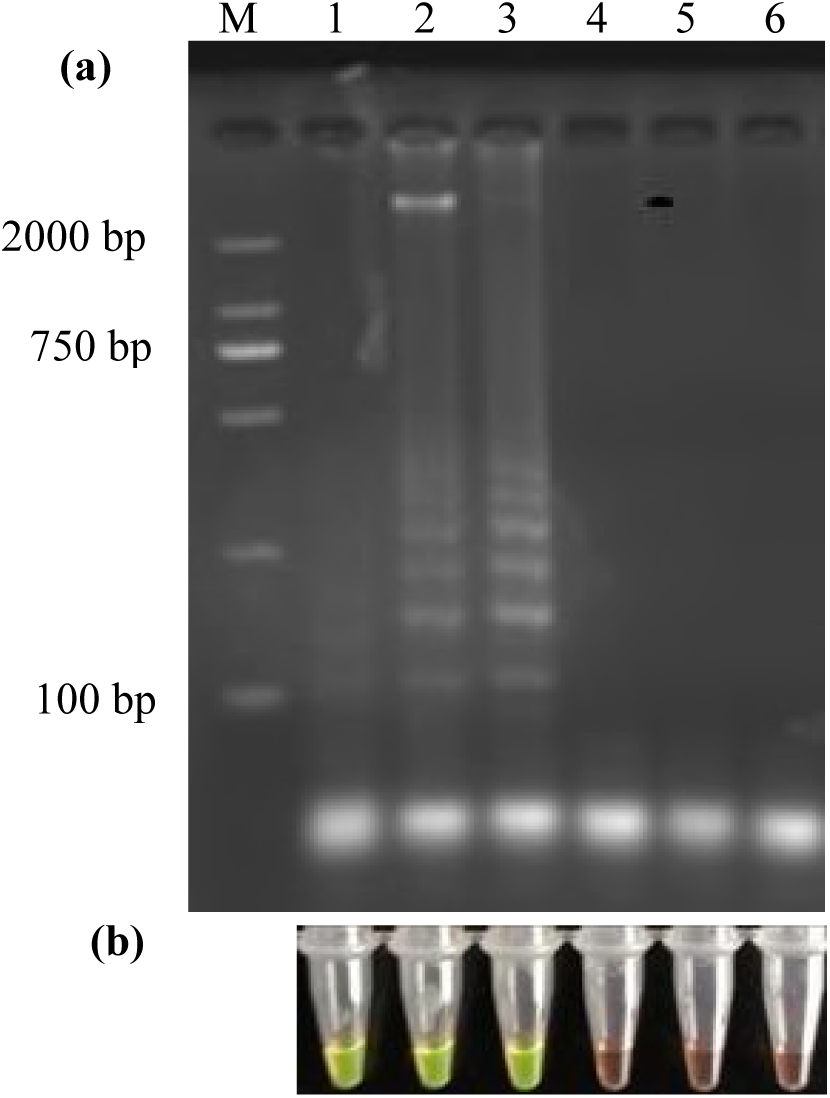
Detection of the *P. aeruginosa nirS* gene with triplicate LAMP assays using *P. aeruginosa* cells as templates. Amplification success was verified by gel electrophoresis (a) and GeneFinder (b). Lanes 1–3 are amplification reactions using 3.36×10^2^ CFU *P. aeruginosa* cells as template; Lanes 4–6 are the negative control (using water as template). The amplification conditions are the same as in Figure 2.

### 3.2 Specificity of LAMP assay

The specificity of the LAMP assay for the detection of *nirS* gene was determined using ∼0.10 ng of genomic DNA from various bacterial species as template (Table 2). Results (Figure S1and Figure S2) indicated the specific amplification of *nirS* from *P. aeruginosa* genomic DNA that contains the cytochrome *cd1*-containing nitrite reductase encoding gene (34). Moreover, no false positive or false negative results were observed when using template DNA from a wide range of bacterial species, indicating good specificity of the LAMP assay for the *nirS* gene.

**Table 2.**
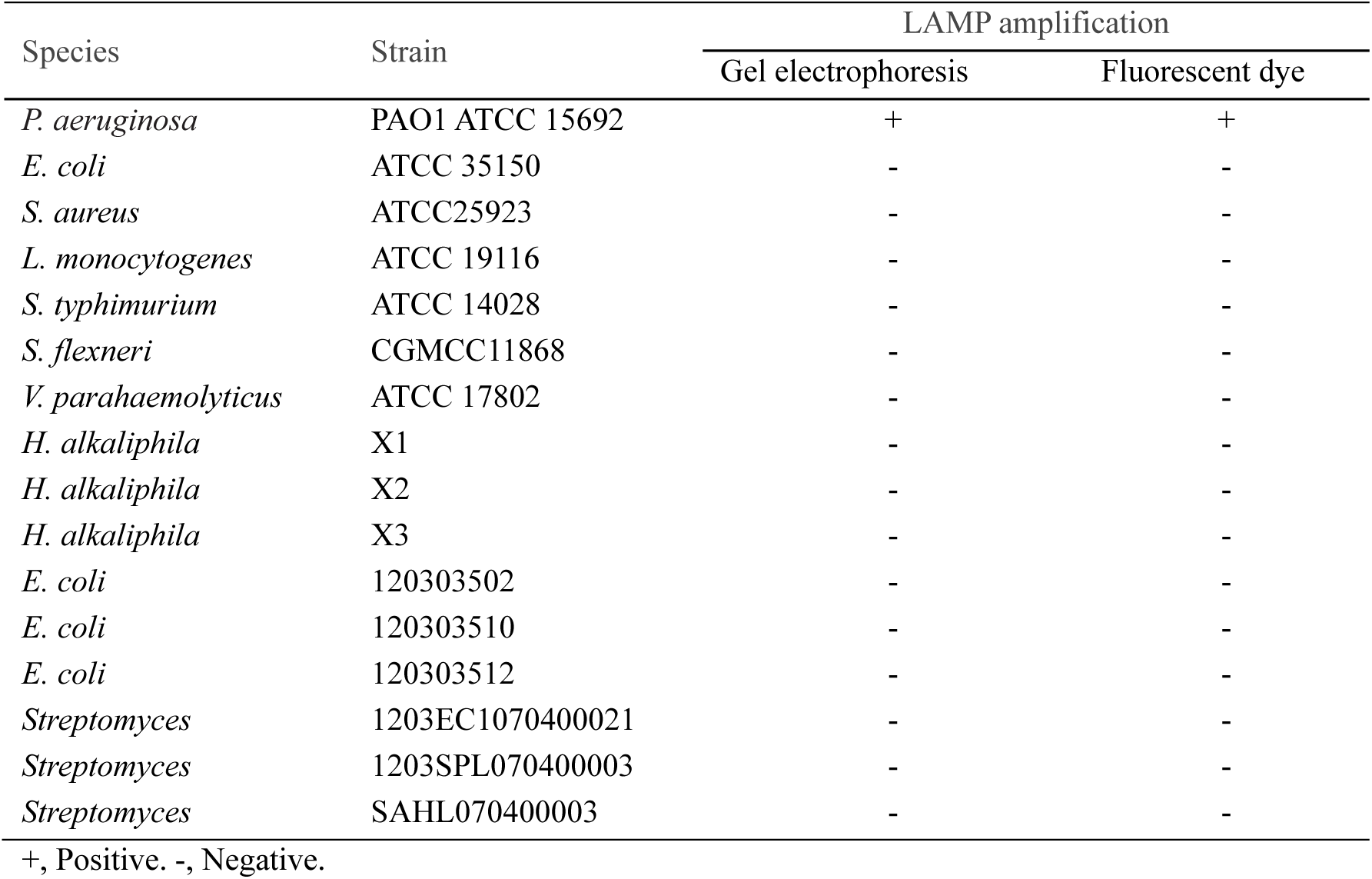
DNA-template based LAMP assays for detecting *nirS* gene of various bacterial species.

Experiments were also conducted to evaluate the specificity of *nirS* detection via cell-template based LAMP assay using ∼10^3^ CFU/reaction from various bacterial species as template. The results from these assays (Table 3) were consistent with those obtained from DNA-template based LAMP assays (Table 2), indicating high specificity of cell-template based amplification under the selected conditions.

**Table 3.**
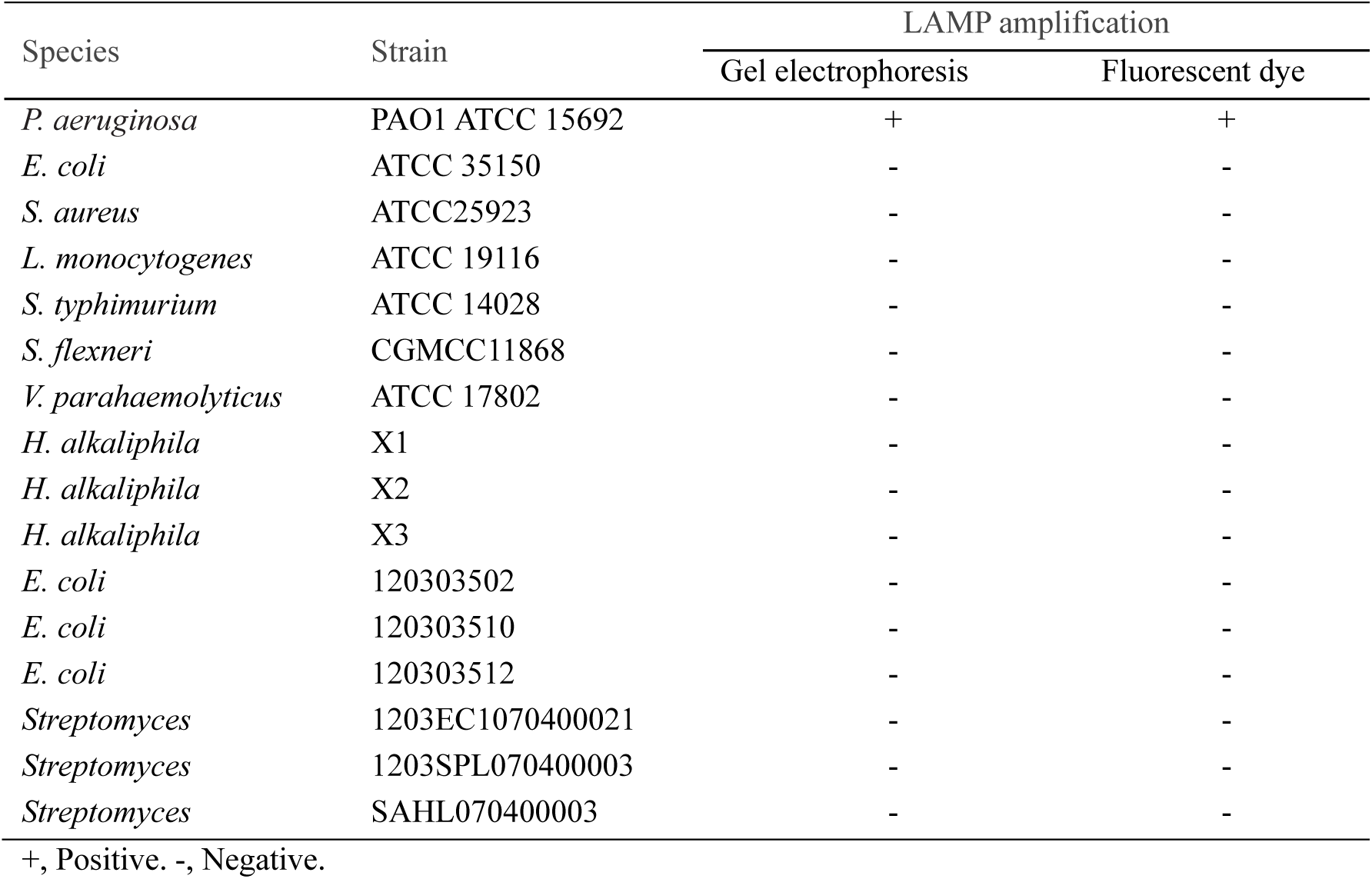
Cell-template based LAMP assays for detecting *nirS* gene of various bacterial species.

### 3.3 Sensitivity of LAMP assay

We assessed the sensitivity of DNA-template based LAMP assay over the amount range of 1.87 fg - 187.00 ng. The results (Figure 4a and 4b) indicated that the LOD was 1.87 pg/reaction with these specified parameters. Below this LOD, no visual detection of amplification products was observed. Moreover, the sensitivity of gel electrophoresis and visual detection were equivalent, suggesting that they were both equally appropriate for determining LAMP amplification success. Using longer incubation times can lower the LOD of LAMP assays at the expense of analysis efficiency (40). Consequently, 60 min was selected as the incubation time for all other reactions. The sensitivity of cell-template based LAMP assays was also evaluated as above with amount of *P. aeruginosa* cells over the range of 3.36 × 10^0^ - 3.36 × 10^8^ CFU/reaction (Figure 5a and 5b). The LOD was 3.36 × 10^2^ CFU/reaction.

**Figure 4.**
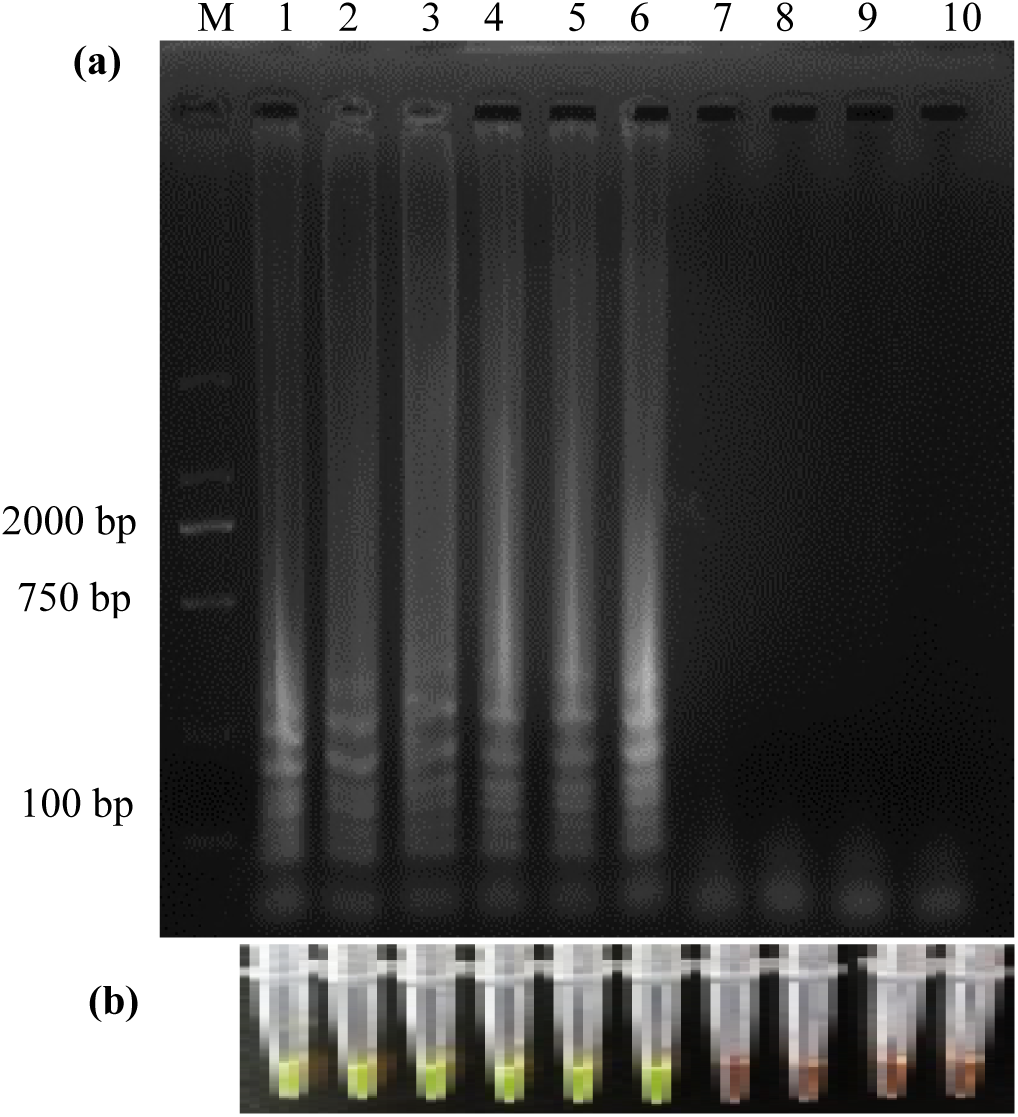
LAMP assay results for *nirS* amplification using varying amounts of *P. aeruginosa* genomic DNA as template. Amplification success was verified by gel electrophoresis (a) and GeneFinder (b). Lanes 1 to 10 correspond to template genomic DNA amounts of 187.00 ng, 18.70 ng, 1.87 ng, 187.00 pg, 18.70 pg, 1.87 pg, 187.00 fg, 18.70 fg, 1.87 fg, and 0.00 fg/reaction, respectively. The amplification conditions are the same as in Figure 2.

**Figure 5.**
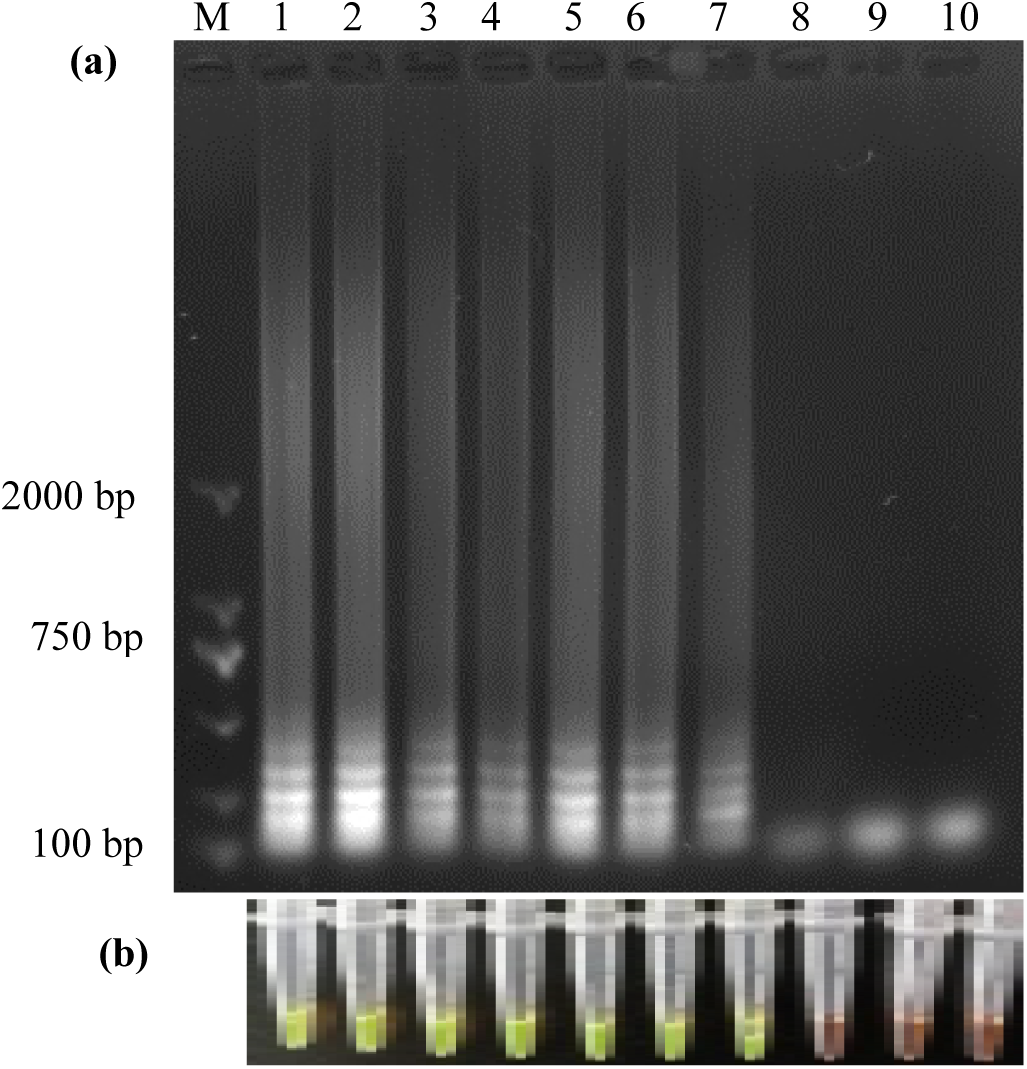
LAMP assay results for *nirS* amplification using varying numbers of *P. aeruginosa* cells as template. Amplification success was verified by gel electrophoresis (a) and GeneFinder (b). Lanes 1 to 10 correspond to template cell numbers of 3.36 × 10^8^ CFU, 3.36 × 10^7^ CFU, 3.36 × 10^6^ CFU, 3.36 × 10^5^CFU, 3.36 × 10^4^ CFU, 3.36 × 10^3^ CFU, 3.36 × 10^2^ CFU, 3.36 × 10^1^ CFU, 3.36 × 10^0^ CFU, and 0.00 CFU, respectively. The amplification conditions are the same as in Figure 2.

### 3.4 Comparison of PCR and LAMP

Using the F3 and B3 primers, experiments were conducted to determine the sensitivity of conventional PCR assay in comparison with the DNA-template based LAMP assay. Genomic DNA amount ranging from 1.87 fg–187.00 ng/reaction were used as template for the reactions. Gel electrophoresis characterization of PCR amplification products indicated no amplification when the DNA template was in a lower amount than 18.70 pg/reaction (Figure 6), but amplification was detected over the range of 18.70 pg to 187.00 ng/reaction. These results indicate a wider dynamic range of the LAMP assays, with 10-fold greater sensitivity than conventional PCR when using genomic DNA. Further, no PCR amplification was detected when *P. aeruginosa* cells were directly added to each PCR reaction mixture over the range of 3.36 × 10^8^ - 3.36 × 10^4^ CFU/reaction.

**Figure 6.**
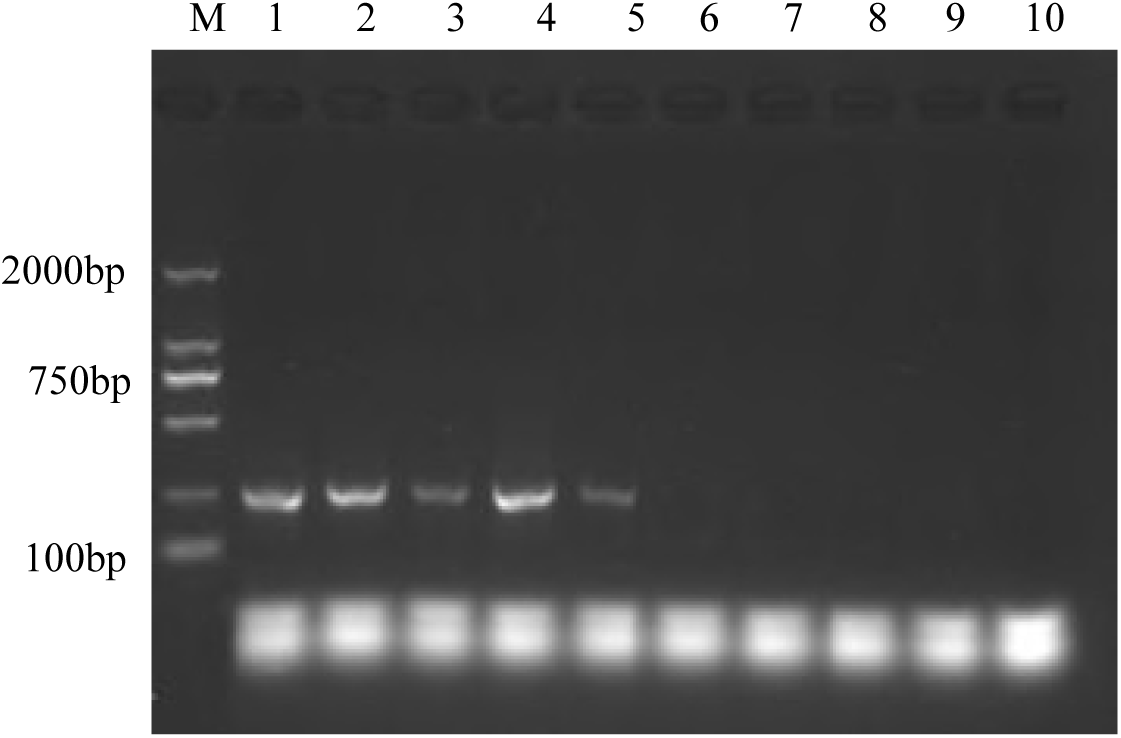
Conventional PCR amplification results for *nirS* using varying amounts of *P. aeruginosa* genomic DNA as template. Amplification success was verified by gel electrophoresis. Lanes 1 to 10 correspond to template genomic DNA amounts of 187.00 ng, 18.70 ng, 1.87 ng, 187.00 pg, 18.70 pg, 1.87 pg, 187.00 fg, 18.70 fg, 1.87 fg, and 0.00 fg/reaction, respectively.

### 3.5 Detection of nirS gene in spiked seawater samples

To investigate the ability of the DNA-template based LAMP and the cell-template based LAMP assays for detecting *nirS* in complex matrices, we spiked seawater samples with *P. aeruginosa* genomic DNA or cells over concentration ranges of 1.27 × 10^2^–1.27 × 10^8^ fg/μL and 1.68×10^1^ - 1.68×10^7^ CFU/mL, respectively. A 2 μL aliquot of the spiked samples was then used as a template in reactions with incubations at 63°C for 60 min. Amplification success was characterized by staining with GeneFinder. Amplifications did not occur with genomic DNA concentrations lower than 1.27×10^4^ fg/μL in the spiked samples (Figure S3). In the cell-template based LAMP assay, a 50 mL mixture of seawater spiked with cells at different concentrations was pretreated by centrifugation to pellet cells. The obtained biomass pellet was then directly used as the template for the cell-template based LAMP assay. Amplifications occurred using every biomass pellet obtained from the spiked samples (Figure S4). When using a 2 μL spiked sample as a template, amplifications only occurred when cell concentrations were greater than 1.68 × 10^4^ CFU/mL (Figure S5).

## 4. Discussion

Denitrification and denitrifying microbial communities have recently received widespread research attention due to their important contributions to the global nitrogen cycle (1,8,41). Functional genes involved in nitrite reduction, especially the cytochrome *cd1*-containing nitrite reductase encoding gene, *nirS*, are commonly used as molecular markers to detect denitrifying populations and potential activities (8,41-44). Concomitantly, the recent development of a novel gene amplification procedure, LAMP, has shown great promise in overcoming the numerous drawbacks of conventional PCR gene amplification methods. In this study, a DNA-template based LAMP assay and a cell-template based LAMP assay were developed to detect *nirS* gene of *P. aeruginosa*. The characteristics of these assays are discussed below and compared against those of conventional PCR assays.

LAMP reactions achieve DNA amplifications using a one-step reaction with a set of target-specific primers (e.g., FIP, BIP, F3, and B3) that recognize six distinct sites flanking the target sequence. The FIP and BIP, each of which contains two functional sequences (one for priming extension in the first stage and the other for self-priming in the second stage) corresponding to the sequences (sense and antisense) of the target dsDNA, play major roles in the LAMP reaction. Catalyzing by *Bst* DNA polymerase with strand displacement activity, LAMP reaction includes two stages. In the first stage, all of four primers are used to start structure-produce. In the second stage, only FIP and BIP are required for realizing cycling amplification. In brief, an ssDNA is released by strand displacement DNA synthesis primed by an F3 and then acts as the template for DNA synthesis primed by both BIP and B3, producing a stem-loop DNA structure. After initiation by one inner primer complementary to the loop on the product, the cycling amplification process is continued by each inner primer alternately. Thus, the specificity is higher than PCR and the final products are stem-loop DNAs with different inverted target repeats and cauliflower-like structures with multiple loops, which are ladder-like patterns characterized by gel electrophoresis (18,20). *NirS* gene is absent in *S. aureus* and *E. coli* genomes, but present in those of *P. aeruginosa* (45), which is consistent with other reports (2, 34). Our amplification results from LAMP specificity assays are consistent with these reports.

PCR activity strongly depends on the cycling of working temperatures, consequently requiring sophisticated equipment to accurately control reaction temperatures. One of the major advantages of the LAMP assay over conventional PCR is eliminating the need for cycling of temperatures, thereby allowing the use of simple, miniature, and affordable amplification devices, in addition to requiring lower energy consumption (21). These features render LAMP assays suitable for use in resource-limited rural areas. Moreover, these advantages make LAMP a promising approach for realizing in-field detection and avoiding cumbersome transportation from sampling sites to specialized laboratories, as is necessary for conventional PCR detection of *nirS* gene from environmental samples (3-5,11-14).

PCR products are typically characterized by gel electrophoresis (5,13,43) or otherwise via quantification with fluorescent probes (5,42,43). In contrast, more quantification approaches can be employed to determine LAMP product amplification, including both endpoint and online patterns. Gel electrophoresis and GeneFinder characterization are both endpoint analyses that are appropriate for LAMP detection, as shown here and elsewhere. In addition, several alternative endpoint methods can be used, including assays with SYBR Green I, Quant-iT PicoGreen, and polyethylenimine, among others. Further, the large amount of white precipitate that is the product of insoluble magnesium pyrophosphate can be used to determine LAMP reaction success with or without centrifugation (21). Online characterization methods can also be used to assess LAMP amplification success including the use of turbidimeters, optical fibers, or spectrophotometers that can monitor LAMP reaction progress based on the formation of magnesium pyrophosphate (21,38). Consequently, the addition of special indicator reagents is unnecessary, further reducing reagent and labor costs. Importantly, instruments for real-time monitoring of LAMP amplification are already commercially available.

The results reported here indicate that conventional PCR assays of *nirS* gene required more than 18.7 fg of template DNA for each reaction, which is consistent with results from Li *et al*. (46). In contrast, the LAMP assay results reported here demonstrate a LOD of 1.87 pg/reaction, indicating a significantly higher sensitivity than conventional PCR, which agrees with previous reports (19,27). Moreover, *nirS* gene detection with conventional PCR assays required cell lysis and subsequent DNA extraction (5,43). Consistent with these observations, we found that PCR amplification could not occur using bacterial cells as the amplification template. DNA extraction, PCR reactions, and electrophoresis typically require >1 h each, and all of these procedures require bulky, specialized equipment. Performing real-time quantitative PCR is much quicker than traditional PCR due to the measurement of reaction results in real time. However, qPCR necessitates expensive probes, even more sophisticated equipment than traditional PCR and is still time consuming. Consequently, conventional and real-time PCR assays are not amenable to detection of *nirS* gene in point-of-care settings. LAMP has the potential to circumvent these problems due to a reduced dependence on pretreatment of samples and the ability to conduct LAMP under isothermal condition (18,21). In particular, the efficacy of cell-template based LAMP assay considerably enhances its application in point-of-care settings (25,30,31). For example, we successfully detected *nirS* gene of *P. aeruginosa* cells over a range of 3.36 × 10^2^ - 3.36 × 10^8^ CFU/reaction. These results further confirm that LAMP assays are less affected by substances that typically inhibit conventional PCR (21,22). Consequently, simpler LAMP assays can be developed by eliminating the DNA extraction step that is necessary prior to conventional PCRs. Further, only 1 h was needed from the addition of template bacterial cells to amplification verification without the need for bulky and sophisticated equipment. Moreover, *nirS* gene of *P. aeruginosa* could be detected in spiked seawater samples with either DNA template or bacterial cells template, further demonstrating the practicality of the LAMP assays, even in complex background matrices. It should be noted, however, that sensitivity of the LAMP assay was clearly affected by the presence of complex co-existing substances in the seawater.

Future investigations of *nirS* amplification via LAMP assays will focus on improving the assays through three target areas. First, the specificity of the LAMP assay towards *nirS* from other organisms including *Pseudomonas aeruginosa* (8), *Pseudomonas stutzeri* (13) and other taxa (46) will be tested to determine its capacity for analyzing denitrifier communities, in general. Second, methods will be developed to eliminate interference from dead cells and extracellular DNA, because only gene expression from viable cells is meaningful towards understanding functional protein expression and consequent denitrification activity. Lastly, a quantitative LAMP assay will be developed to determine the relationship between *nirS* gene copy abundance in viable microbial cells and denitrifying efficiency.

## 5. Conclusions

Herein, LAMP assays were developed for detecting cytochrome *cd1*-containing nitrite reductase encoding gene, *nirS*, for the first time. The developed primer sets recognized the target sequence with high specificity, as characterized by gel electrophoresis or GeneFinder visualization. Optimized incubation temperature at 63°C and a time of 60 min led to an obtained LOD of 1.87 pg. Importantly, this limit was lower than that of conventional PCR assays by an order of magnitude.

One of the unique advantages of LAMP amplification is the capacity to amplify DNA directly from bacterial cell templates, owing to its higher resistance to inhibition than conventional PCR assays. The cell-template based LAMP also exhibited high specificity. After optimizing amplification conditions, a LOD of 3.36 × 10^2^ CFU/reaction was obtained. Moreover, only 1 h was needed from the addition of bacterial cells to the confirmation of *nirS* amplification, and bulky and sophisticated equipment were not required.

Overall, the LAMP assay presented here was superior to conventional PCR assays in terms of sensitivity, specificity, turnover-time, simplicity, and cost. Importantly, the LAMP assays described here are ready to use for in-field applications. Their practicality using environmental samples was preliminarily demonstrated using seawater samples spiked with genomic DNA or *P. aeruginosa* cells. Additional studies are planned to further improve the accuracy and general applicability of the LAMP assays reported here.

## Acknowledgements

This work was supported by the National Key R&D Program of China (YS2017YFGH000215), the Special Scientific Research Funds for Central Non-profit Institutes, the Yellow Sea Fisheries Research Institute, the Chinese Academy of Fishery Sciences (20603022018020, 20603022016003), and the National Key R&D Program of China (2017YFE1015200).

---

**Scheme 1.**
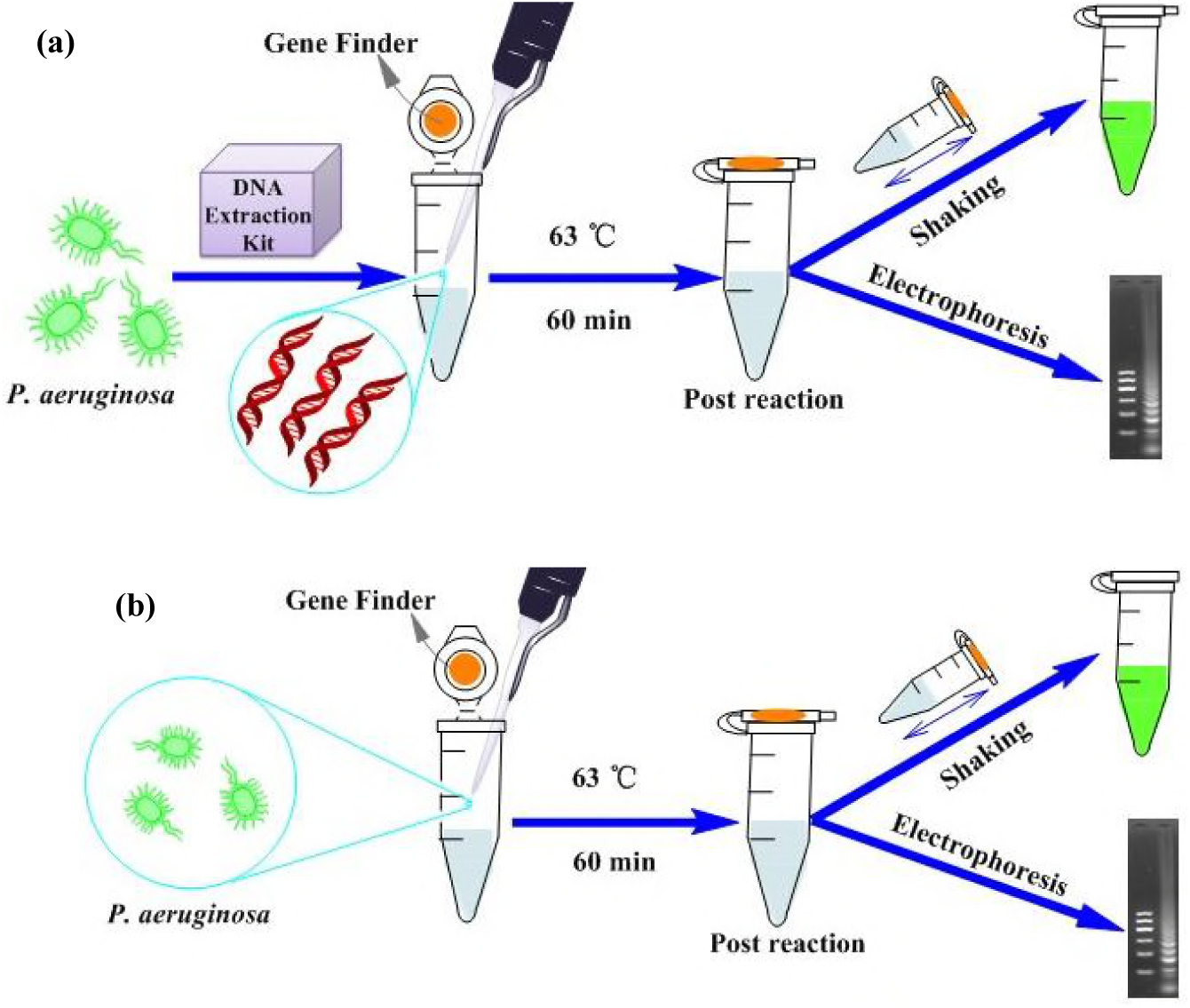
Methodological schematics for the DNA-template based LAMP assay (a) and the cell-template based direct LAMP assay (b) for detecting the *nirS* genes of *P. aeruginosa*.

## Reference

1. Gruber N, Galloway JN. 2008. An Earth-system perspective of the global nitrogen cycle. Nature 451:293–296.

2. Throback IN, Enwall K, Jarvis A, et al. 2010. Reassessing PCR primers targeting nirS, nirK and nosZ genes for community surveys of denitrifying bacteria with DGGE. Fems Microbiology Ecology 49(3):401–417.

3. Braker G, Zhou J, Wu L, et al. 2000. Nitrite reductase genes (nirK and nirS) as functional markers to investigate diversity of denitrifying bacteria in pacific northwest marine sediment communities. Applied and Environmental Microbiology 66(5):2096–2104.

4. Francis CA, O’Mullan GD, Cornwell JC, Ward BB. 2013. Transitions in nirS-type denitrifier diversity, community composition, and biogeochemical activity along the Chesapeake Bay estuary. Frontiers in Microbiology 4:237–237.

5. Zheng Y, Hou L, Liu M, Gao J, Yin G, Li X, Deng D, Lin X, Jiang X, Chen F, Zong H, Zhou J. 2015. Diversity, Abundance, and Distribution of nirS-Harboring Denitrifiers in Intertidal Sediments of the Yangtze Estuary. Microbial Ecology 70:30–40.

6. Zumft WG. 1997. Cell biology and molecular basis of denitrification. Microbiology and Molecular Biology Reviews 61:533–616.

7. Pratscher J, Stichternoth C, Fichtl K, Schleifer K, Braker G. 2009. Application of recognition of individual genes-fluorescence in situ hybridization (RING-FISH) to detect nitrite reductase genes (nirK) of denitrifiers in pure cultures and environmental samples. Applied and Environmental Microbiology 75:802.

8. Heylen K, Gevers D, Vanparys B, Wittebolle L, Geets J, Boon N, Vos PD. 2006. The incidence of nirS and nirK and their genetic heterogeneity in cultivated denitrifiers. Environmental Microbiology 8(11):2012–2021.

9. Yoshida M, Ishii S, Otsuka S. 2010. nirK-harboring denitrifiers are more responsive to denitrificationinducing conditions in rice paddy soil than nirS-harboring bacteria. Microbes and Environments 25(1):45–48.

10. Jones CM, Stres B, Rosenquist M, Hallin S. 2008. Phylogenetic analysis of nitrite, nitric oxide, and nitrous oxide respiratory enzymes reveal a complex evolutionary history for denitrification. Molecular Biology and Evolution 25(9):1955–1966.

11. Yang Y, Zhao J, Jiang Y, Hu Y, Zhang M, Zeng Z. 2017. Response of bacteria harboring nirS and nirK genes to different N fertilization rates in an alkaline northern Chinese soil. European Journal of Soil Biology 82:1–9.

12. Priemé A, Braker G, Tiedje M. 2002. Diversity of nitrite reductase (nirK and nirS) gene fragments in forested upland and wetland soils. Applied and Environmental Microbiology 68(4):1893–1900.

13. Braker G, Fesefeldt A, Witzel KP. 1998. Development of PCR primer systems for amplification of nitrite reductase genes (nirK and nirS) to detect denitrifying bacteria in environmental samples. Applied and Environmental Microbiology 64(10):3769–3775.

14. Harbi B, Chaieb K, Jabeur C, Bakhrouf A. 2010. PCR detection of nitrite reductase genes (nirK and nirS) and use of active consortia of constructed ternary adherent staphylococcal cultures via mixture design for a denitrification process. World Journal of Microbiology and Biotechnology 26(3):473–480.

15. Jayakumar A, OMullan GD, Naqvi SWA, Ward BB. 2009. Denitrifying bacterial community composition changes associated with stages of denitrification in oxygen minimum zones. Microbial Ecology 58(2):350–362.

16. Simth J, Wagner-Riddle C, Dunfield K. 2010. Season and management related changes in the diversity of nitrifying and denitrifying bacteria over winter and spring. Applied Soil Ecology 44(2):138–146.

17. Taroncher-Oldenburg G, Griner EM, Francis CA, Ward BB. 2003. Oligonucleotide microarray for the study of functional gene diversity in the nitrogen cycle in the environmental. Applied and Environmental Microbiology 69(2):1159–1171.

18. Zhao Y, Chen F, Li, Q, Wang L, Fan C. 2015. Isothermal amplification of nucleic acids. Chemical Reviews 115:12491–12545.

19. Verma S, Singh R, Sharma V, Avtar-Bumb R, Singh-Negi N, Ramesh V, Salotra P. 2017. Development of a rapid loop-mediated isothermal amplification assay for diagnosis and assessment of cure of Leishmania infection. BMC Infectious Diseases 17(1):223.

20. Notomi T, Okayama H, Masubuchi H, Yonekawa T, Watanabe K. 2000. Loop-mediated isothermal amplification of DNA. Nucleic Acids Research 28(12):e63.

21. Zhang X, Lowe SB, Gooding JJ. 2014. Brief review of monitoring methods for loop-mediated isothermal amplification (LAMP). Biosensors and Bioelectronics 61:491–499.

22. Abdul-Ghani R, Al-Mekhlafi AM, Karanis P. 2012. Loop-mediated isothermal amplification (LAMP) for malarial parasites of humans: Would it come to clinical reality as a point-of-care test?. Acta Tropica 122(3):233–240.

23. Safavieh M, Kanakasabapathy MK, Tarlan F, Ahmed MU, Zourob M. 2016. Emerging loop-mediated isothermal amplification-based microchip and microdevice technologies for nucleic acid detection. Acs Biomaterials Science and Engineering 2(3):278.

24. Njiru ZK. 2012. Loop-mediated isothermal amplification technology: Towards point of care diagnostics. PLoS Neglected Tropical Diseases 6(6):e1572.

25. Williams MR, Stedtfeld RD, Waseem H, Stedtfeld T, Upham B. 2017. Implications of direct amplification for measuring antimicrobial resistance using point-of-care devices. Analytical Methods 9:(8).

26. Lee D, Kim YT, Lee JW, Kim DH, Seo TS. 2016. An integrated direct loop-mediated isothermal amplification microdevice incorporated with an immunochromatographic strip for bacteria detection in human whole blood and milk without a sample preparation step. Biosensors and Bioelectronics 79:273–279.

27. Koizumi N, Nakajima C, Harunari T, Tanikawa T, Tokiwa T. 2012. A new loop-mediated isothermal amplification method for rapid, simple, and sensitive detection of Leptospira spp. in urine. Journal of Clinical Microbiology 50(6):2072–2074.

28. Soejima M, Egashira K, Kawano H, Kawaguchi A, Sagawa K. 2011. Rapid detection of haptoglobin gene deletion in alkaline-denatured blood by loop-mediated isothermal amplification reaction. Journal of Molecular Diagnostics 13(3):334–339.

29. Kim MJ, Kim HY. 2018. Direct duplex real-time loop mediated isothermal amplification assay for the simultaneous detection of cow and goat species origin of milk and yogurt products for field use. Food Chemistry 246:26–31.

30. Kanitkar YH, Stedtfeld RD, Hatzinger PB, Hashsham SA, Cupples AM. 2017. Development and application of a rapid, user-friendly, and inexpensive method to detect Dehalococcoides sp. reductive dehalogenase genes from groundwater. Applied Microbiology and Biotechnology:1-9.

31. Stedtfeld RD, Stedtfeld TM, Samhan F, Kanitkar YH,Hatzinger PB. 2016. Direct loop mediated isothermal amplification on filters for quantification of Dehalobacter in groundwater. Journal of Microbiological Methods 131:61–67.

32. Zhang X, Qu K, Li Q, Cui Z, Zhao J, Sun X. 2011. Recording the reaction process of loop-mediated isothermal amplification (LAMP) by monitoring the voltammetric response of 2’-deoxyguanosine 5’-triphosphate. Electroanalysis 23:2438–2445.

33. Zhang X, Liu W, Lu X, Justin GJ, Li Q, Qu K. 2014. Monitoring the progression of loop-mediated isothermal amplification (LAMP) using conductivity. Analytical Biochemistry 466:16–18.

34. Calmels S, Ohshima H, Henry Y, Henry Y, Bartsch H. 1996. Characterization of bacterial cytochrome cd1-nitrite reductase as one enzyme responsible for catalysis of nitrosation of secondary amines. Biomedical and General Engineering:392-402.

35. Lin HL, Lin CC, Lin YJ, Lin HC, Shih CM, Chen CR, Huang RN, Kuo TC. 2010. Revisiting with a relative-density calibration approach the determination of growth rates of microorganisms by use of optical density data from liquid cultures. Applied and Environmental Microbiology 76(5):1683–1685.

36. Zhang X, Jiang X, Yang Q, Wang X, Zhang Y, Zhao J, Qu K, Zhao C. 2018. Online monitoring of bacterial growth with electrical sensor. Analytical Chemistry 90(10):6006–6011.

37. Feng J, Dai Z, Tian X, Jiang X. 2108. Detection of Listeria monocytogenes based on combined aptamers magnetic capture and loop-mediated isothermal amplification. Food Control 85: 443–452.

38. Tomita N, Mori Y, Kanda H, Notomi T. 2008. Loop-mediated isothermal amplification (LAMP) of gene sequences and simple visual detection of products. Nature Protocols 3(5):877–882.

39. Balbin MM, Belotindos LP, Abes NS, Mingala CN. 2014. Caprine arthritis encephalitis virus detection in blood by loop-mediated isothermal amplification (LAMP) assay targeting the proviral gag region. Diagnostic Microbiology and Infectious Disease 79(1):37–42.

40. Mori Y, Kitao M, Tomita N, Notomi T. 2004. Real-time turbidimetry of LAMP reaction for quantifying template DNA. Journal of Biochemical and Biophysical Methods 59(2):145–157.

41. Chen Y, Zhou W, Li Y, Zhang J, Zeng G, Huang A, Huang J. 2014. Nitrite reductase genes as functional markers to investigate diversity of denitrifying bacteria during agricultural waste composting. Applied Microbiology and Biotechnology 98(9):4233–4243.

42. Gomes J, Khandeparker R, Bandekar M, Meena RM, Ramaiah N. 2017. Quantitative analyses of denitrifying bacterial diversity from a seasonally hypoxic monsoon governed tropical coastal region. Deep Sea Research Part II Topical Studies in Oceanography.

43. Gao J, Hou L, Zheng Y, Liu M, Yin G, Li X, Lin X, Yu C, Wang R, Jiang X, Sun X. 2016. nirS-Encoding denitrifier community composition, distribution, and abundance along the coastal wetlands of China. Applied Microbiology and Biotechnology100(19):1–10.

44. Li M, Hong Y, Cao H, Gu J. 2013. Community structures and distribution of anaerobic ammonium oxidizing and nirs-encoding nitrite-reducing bacteria in surface sediments of the south china sea. Microbial Ecology 66(2):281–296.

45. Hallin S, Lindgren PE. 1999. PCR detection of genes encoding nitrite reductase in denitrifying bacteria. Applied and environmental microbiology 65(4):1652–1657.

46. Li M, Ford T, Li X, Gu J. 2011. Cytochrome cd1-containing nitrite reductase encoding gene nirs as a new functional biomarker for detection of anaerobic ammonium oxidizing (anammox) bacteria. Environmental Science and Technology 45(8):3547.

